# Codon optimization, not gene content, predicts *XYL*ose metabolism in budding yeasts

**DOI:** 10.1101/2022.06.10.495693

**Authors:** Rishitha L. Nalabothu, Kaitlin J. Fisher, Abigail Leavitt LaBella, Taylor A. Meyer, Dana A. Opulente, John F. Wolters, Antonis Rokas, Chris Todd Hittinger

## Abstract

Xylose is the second most abundant monomeric sugar in plant biomass. Consequently, xylose catabolism is an ecologically important trait for saprotrophic organisms, as well as a fundamentally important trait for industries that hope to convert plant mass to renewable fuels and other bioproducts using microbial metabolism. Although common across fungi, xylose catabolism is rare within Saccharomycotina, the subphylum that contains most industrially relevant fermentative yeast species. Several yeasts unable to consume xylose have been previously reported to possess complete predicted xylolytic metabolic pathways, suggesting the absence of a gene-trait correlation for xylose metabolism. Here, we measured growth on xylose and systematically identify *XYL* pathway orthologs across the genomes of 332 budding yeast species. We found that most yeast species possess complete predicted xylolytic pathways, but pathway presence did not correlate with xylose catabolism. We then quantified codon usage bias of *XYL* genes and found that codon optimization was higher in species able to consume xylose. Finally, we showed that codon optimization of *XYL2*, which encodes xylitol dehydrogenase, positively correlated with growth rates in xylose medium. We conclude that gene content cannot predict xylose metabolism; instead, codon optimization is now the best predictor of xylose metabolism from yeast genome sequence data.

**Significance Statement:** In the genomic era, strategies are needed for the prediction of metabolic traits from genomic data. Xylose metabolism is an industrially important trait, but it is not found in most yeast species heavily used in industry. Because xylose metabolism appears rare across budding yeasts, we sought to identify a computational means of predicting which species are capable of xylose catabolism. We did not find a relationship between gene content and xylose metabolism traits. Rather, we found that codon optimization of xylolytic genes was higher in species that can metabolize xylose, and that optimization of one specific gene correlated with xylose-specific growth rates. Thus, codon optimization is currently the only means of accurately predicting xylose metabolism from genome sequence data.

## Introduction

Xylose is the most abundant pentose sugar and the second most abundant monomeric sugar in plant biomass, second only to glucose. Xylose occurs in xylan polymers in hemicellulose; therefore, the ability to hydrolyze xylan and oxidize xylose for energy is a common trait in saprophytic fungi (1). Metabolic conversion of xylose is also a critical process in the efficient conversion of lignocellulosic biomass into biofuels and other bioproducts via fermentation by industrially leveraged yeast species. Unlike filamentous fungi, native xylose assimilation appears to be a somewhat rare trait within budding yeasts. *Saccharomyces cerevisiae* is the choice microbe for the industrial production of the vast majority of biofuels due to its high ethanol tolerance, high glycolytic and fermentative capacity, and amenability to genetic engineering (2). However, *S. cerevisiae* requires genetic engineering to metabolize xylose, and even engineered strains are often inefficient in the fermentation of lignocellulosic xylose (3–6). This has led to the suggestion that cost-effective industrial conversion of xylose would be better achieved using native pentose-fermenting yeast species. One successful approach to identifying xylolytic species is the isolation of yeasts from xylose-rich environments, such as rotting logs and the guts of wood-boring beetles (7–9). Given that budding yeast genomes are increasingly available (10,11), a simpler means of identifying xylolytic yeasts through genome data would facilitate the discovery of additional xylose-metabolizing yeasts.

The budding yeast xylose catabolism pathway was first described in *Cyberlindnera jadinii* and *Candida albicans* (12–14), and most subsequent characterization has focused on xylose-fermenting genera, including *Scheffersomyces* and, more recently, *Spathaspora* (15–17). The native enzymatic pathway consists of three genes: *XYL1, XYL2*, and *XYL3. XYL1* and *XYL2* encode a xylose reductase (XR) and xylitol dehydrogenase (XDH), respectively, which function in the oxidoreductive conversion of xylose to xylulose with xylitol as an intermediate. *XYL3* encodes a xylulokinase (XKS), which phosphorylates xylulose to xylulose-5-phosphate to be fed into the non-oxidative branch of the pentose phosphate pathway. The identification of yeasts with complete pathways that were nonetheless unable to grow on xylose in previous surveys suggests a weak or absent gene-trait association between complete *XYL* pathways and xylose assimilation traits (11,18).

In addition to a complete *XYL* pathway, other genetic and regulatory features may be important in determining xylose metabolic traits. Most studies have focused on the role of redox imbalance, which is thought to be produced by the different cofactor preferences of XR and XDH, which prefer NADPH and NAD^+^, respectively (19). This hypothesis is supported by the observation that some well-studied yeasts that efficiently metabolize xylose have evolved XR enzymes able to use NADH in addition to or in lieu of NADPH (17,20,21).Recently, it has been suggested that changes to cofactor preference in methylglyoxal reductase (encoded by *GRE2*) may also alleviate redox imbalance in xylo-fermentative yeasts (22). Additional properties, such as transporter presence or copy number and the expression of other metabolic genes, have also been implicated in xylose utilization (18). It is difficult to say how broadly applicable any of these explanations may be because the presence of *XYL* genes in the absence of xylose catabolism has only been studied in a handful of related yeast species. Thus, we do not know the extent of this lack of association across budding yeasts and whether other genome characteristics would better predict xylose metabolism.

The identification of some yeasts with complete *XYL* pathways that lack xylose assimilation suggests that xylose utilization may be much more difficult to predict based on gene content than many other metabolic traits, such as galactose utilization (10,11). An alternative strategy to predicting metabolic traits from gene content is evaluating specific metabolic genes for evidence of selection. Measuring selection on codon usage is one such approach. Among metrics developed to measure codon usage bias (23–25), codon optimization captures how well matched individual codons are to their respective tRNA copy numbers in a given genome (26). Accordingly, a codon with a low-copy corresponding tRNA is less optimized than a codon with a high-copy corresponding tRNA. The codon optimization index of a gene therefore measures the concordance between its transcript and the cellular tRNA pool and has repeatedly been shown to correlate with gene expression levels (27–29). Recent work has shown that codon usage is under translational selection in most fungal species (30), including within budding yeasts (31). Studies examining the relationship between codon usage and metabolism in fungi have found that codon bias is elevated in genes encoding important metabolic pathways (32), and further, that codon optimization of metabolic genes is predictive of growth in corresponding conditions (33). Codon optimization of xylolytic genes has not been studied, but we hypothesize that it may be more useful than gene content in predicting which budding yeast species are well-adapted to xylose metabolism.

Here, we measure growth on xylose and systematically identify *XYL* pathway orthologs across 332 publicly available budding yeast genomes (10). In agreement with previous work, we find that an intact *XYL* pathway often does not confer xylose assimilation. We then generate codon optimization indices for all *XYL* homologs and show that, for all three *XYL* genes, codon optimization is significantly higher in species that can consume xylose than in species that cannot. We also demonstrate that kinetic growth rates on xylose are significantly positively correlated with codon optimization of *XYL2*, which underscores the importance of *XYL2* expression levels in xylose metabolism. Collectively, our study identifies a novel means of predicting xylolytic traits from genome sequence alone.

## Results

### Identification *of XYL* homologs across 332 budding yeast species

We were able to detect at least one of the three *XYL* pathway genes in 325 of 332 species (Figure 1). Complete pathways were found in 270 species. We were unable to detect any *XYL* genes in seven species. Six of the seven species with no detected *XYL* homologs were the six representative species of the *Wickerhamiella*/*Starmerella* (W/S) clade, so it appears that the entire *XYL* pathway has been lost in this clade. *XYL1* and *XYL2* have evidence of gene duplications, losses, transfers, and multiple origins prior to the origin of Saccharomycotina, as well as within the budding yeasts. However, due to the sheer breadth of evolutionary distance in this group, confident elucidation of the complete gene history for these genes is intractable with current taxon sampling.

**Figure 1.**
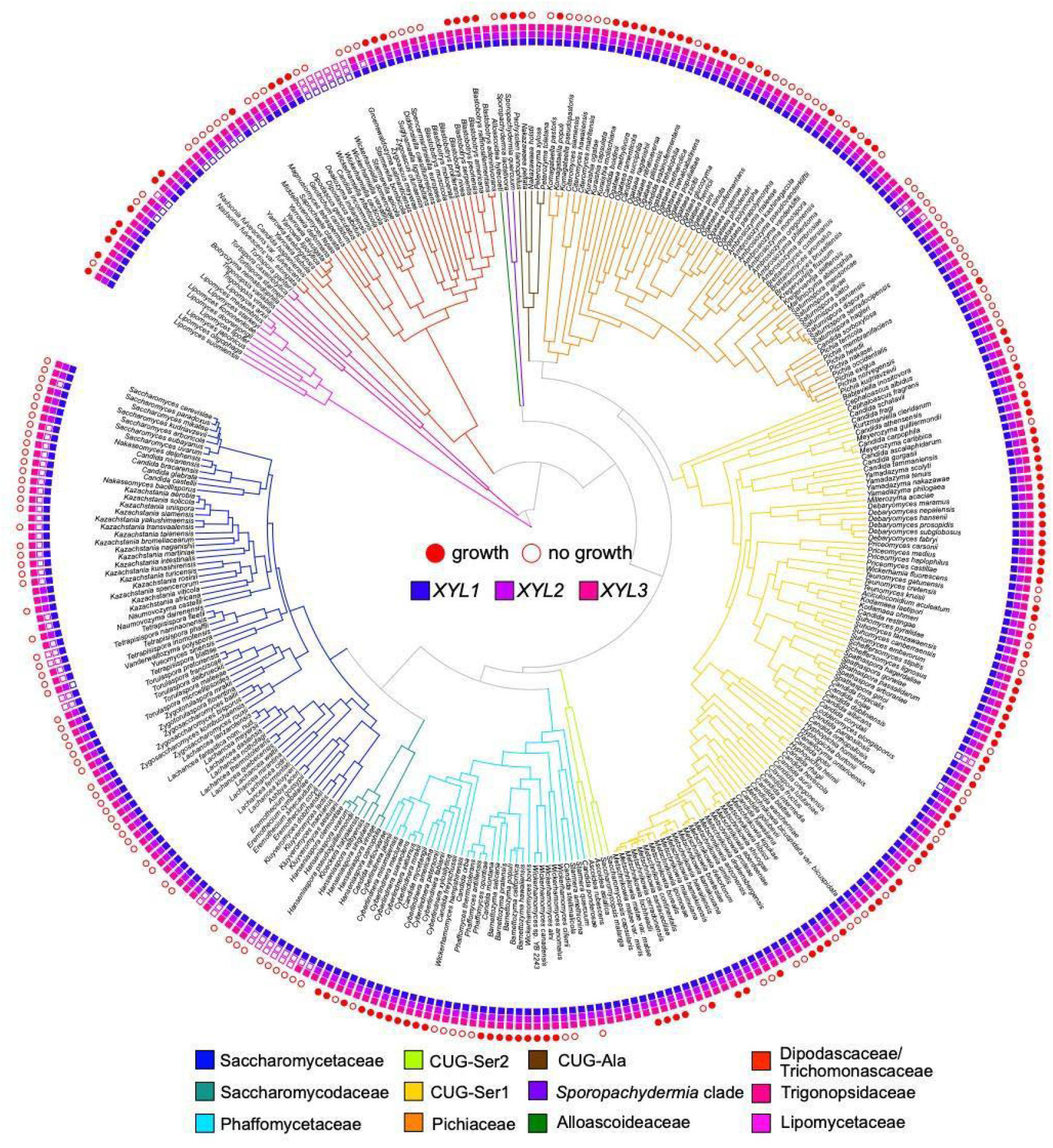
*XYL* pathway presence and xylose growth across 332 Saccharomycotina species. Major yeast clades are depicted by branch color. Presence of *XYL* homologs is indicated by filled boxes. Complete pathways of *XYL1, XYL2*, and *XYL3* were found in 270 species. Species with non-zero growth rates in xylose medium are indicated by a filled red circle, and species unable to assimilate xylose are indicated by an empty red circle. Species without circles were not assayed for growth. Time-calibrated phylogeny from (10).

The phylogenies of *XYL1* and *XYL2* homologs were able to resolve previously ambiguous *S. cerevisiae* orthology (Figures S4-S6). *GRE3* has known xylose reductase activity, but it has been annotated as a nonspecific aldo-keto reductase and believed to be distinct from the XR-encoding genes of xylose-fermenting yeasts (34–36). We found definitive phylogenetic evidence that *GRE3* is a member of the xylose-reductase gene family that is orthologous to the *XYL1* genes of more distantly related yeasts (Fig. S4). In contrast, *S. cerevisiae* is known to contain a *XYL2* homolog, but the function of *XYL2* has remained unclear given the inability of most *S. cerevisiae* strains to metabolize xylose. The nearly identical *S. cerevisiae* paralogs *SOR1* and *SOR2* also fell within the *XYL2* clade of the family Saccharomycetaceae. *SOR1* and *SOR2* are annotated as encoding sorbitol dehydrogenases and are upregulated in response to sorbose and xylose (36) (Fig. S5).

The *XYL2* gene phylogeny showed more evidence of gene diversification and retention than was expected, given that species of the family Saccharomycetaceae are generally not able to use xylose as a carbon source. To further clarify *XYL2* evolution within the Saccharomycetaceae, we generated a maximum likelihood tree of the *XYL2* homologs within the Saccharomycetaceae and included *S. cerevisiae XDH1*, a gene encoding a xylitol dehydrogenase present in some wine strains (but not the S288C reference strain) that was previously identified as being sufficient for weak xylose utilization (37). The resulting tree supports an ancestral duplication of *XYL2*, which produced two distinct paralogous lineages that we name the *SOR* lineage and the *XYL2* lineage based on the *S. cerevisiae* paralogs contained therein (Figure S6). The *XYL2* lineage homolog was preferentially retained by most Saccharomycetaceae species, while a handful retained only the *SOR* paralog, and a few retained both. The tree also supported a few subsequent duplications, including the lineage-specific duplication of *SOR1/SOR2* in *S. cerevisiae*. The phylogeny also showed that the *XDH1* gene identified in Wenger et al. (37) is orthologous to *S. cerevisiae SOR1/SOR2*, not to *S. cerevisiae XYL2*. The protein sequence is also identical to the *Torulaspora microellipsoides SOR* homolog, further corroborating a known 65kb transfer from *T. microellipsoides* to the *S. cerevisiae* EC1118 wine strain and its relatives (38).

### Neither pathway completeness nor copy number predicts growth

Previous surveys of an order of magnitude fewer taxa have suggested a weak or absent gene-trait association between complete *XYL* pathways and xylose assimilation traits (11,17,18). To rigorously test the relationship between gene content and xylose consumption, we measured maximum growth rates in a minimal medium containing xylose as the sole carbon source for 281 of the 332 species examined, which included 236 species found to have complete *XYL* pathways. We found only one species lacking a complete *XYL* pathway, *Candida sojae*, which was able to grow on xylose. Although no *XYL* genes were detected in *C. sojae*, the discrepancy with growth is likely because the *C. sojae* genome assembly queried is incomplete (10). To determine the extent to which pathway presence predicts growth, we examined quantitative growth data in minimal medium containing xylose as the sole carbon source for 236 of the 270 species with complete pathways. We found that a complete *XYL* pathway only conferred a 52% probability of exhibiting a non-zero growth in xylose medium (Fig. 2A), and that 113 species with complete pathways were nonetheless unable to assimilate xylose. We also found that *XYL1* and *XYL*2 copy number did not predict growth, as species with multiple copies of *XYL1* or *XYL2* genes were no more likely to grow than single-copy species (*XYL1* p=0.52, *XYL2* p=0.49, two-tailed Fisher’s exact, Fig. 2B-C).

**Figure 2.**
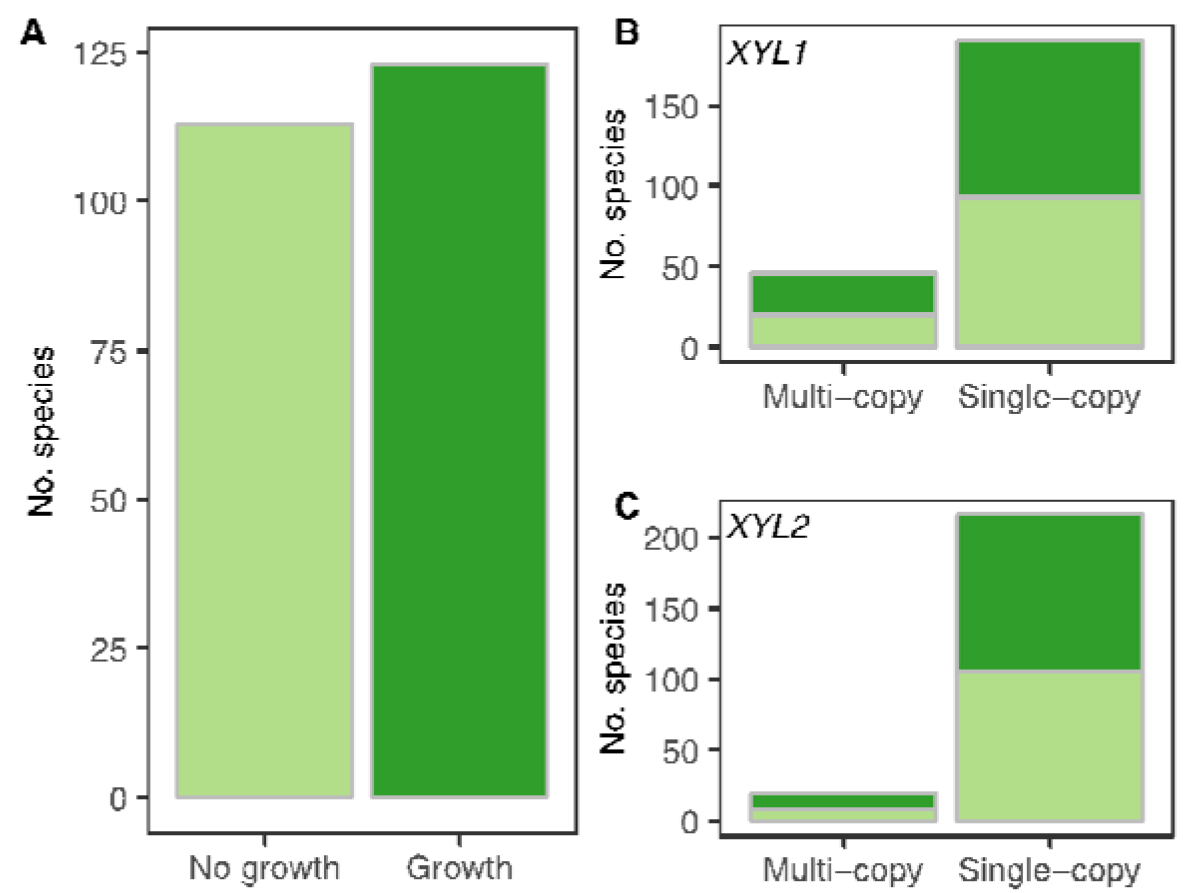
Neither pathway completeness nor copy number predicts xylose metabolism. A) Of the 236 species with complete pathways assayed for growth, 52% (123 species) were able to grow with xylose as a sole carbon source (dark green), while 48% (113 species) did not grow in xylose medium (light green). B-C) *XYL1* and *XYL2* were frequently found to be multi-copy. Two-tailed Fisher’s exact test conducted to determine the impact of multiple copies of *XYL1* and *XYL2* on growth. Dark green bars represent the number of species able to grow, and light green bars represent species unable to grow. Species with multiple copies of either gene were no more likely to grow than species with single copies of *XYL1* or *XYL2* (*XYL1* p=0.52, *XYL2* p=0.49, two-tailed Fisher’s exact test).

### *XYL*1 and *XYL*2 are highly codon-optimized

We next examined codon optimization of the *XYL* pathway genes to determine if codon optimization indices would be more useful in predicting metabolic capabilities than *XYL* gene presence. Codon optimization indices (estAI values) of *XYL* pathway homologs were calculated for 320 of the 325 species in which a *XYL1, XYL2*, or *XYL3* gene was detected. *XYL1* and *XYL2* estAI distributions were both heavily skewed with median estAI values of 0.94 and 0.83, values that equate to a higher optimization than 94% and 83% of the coding genome of an individual species, respectively. *XYL3* estAI values were more variable with a lower median optimization index of 0.55 (Fig. 3A).

**Figure 3.**
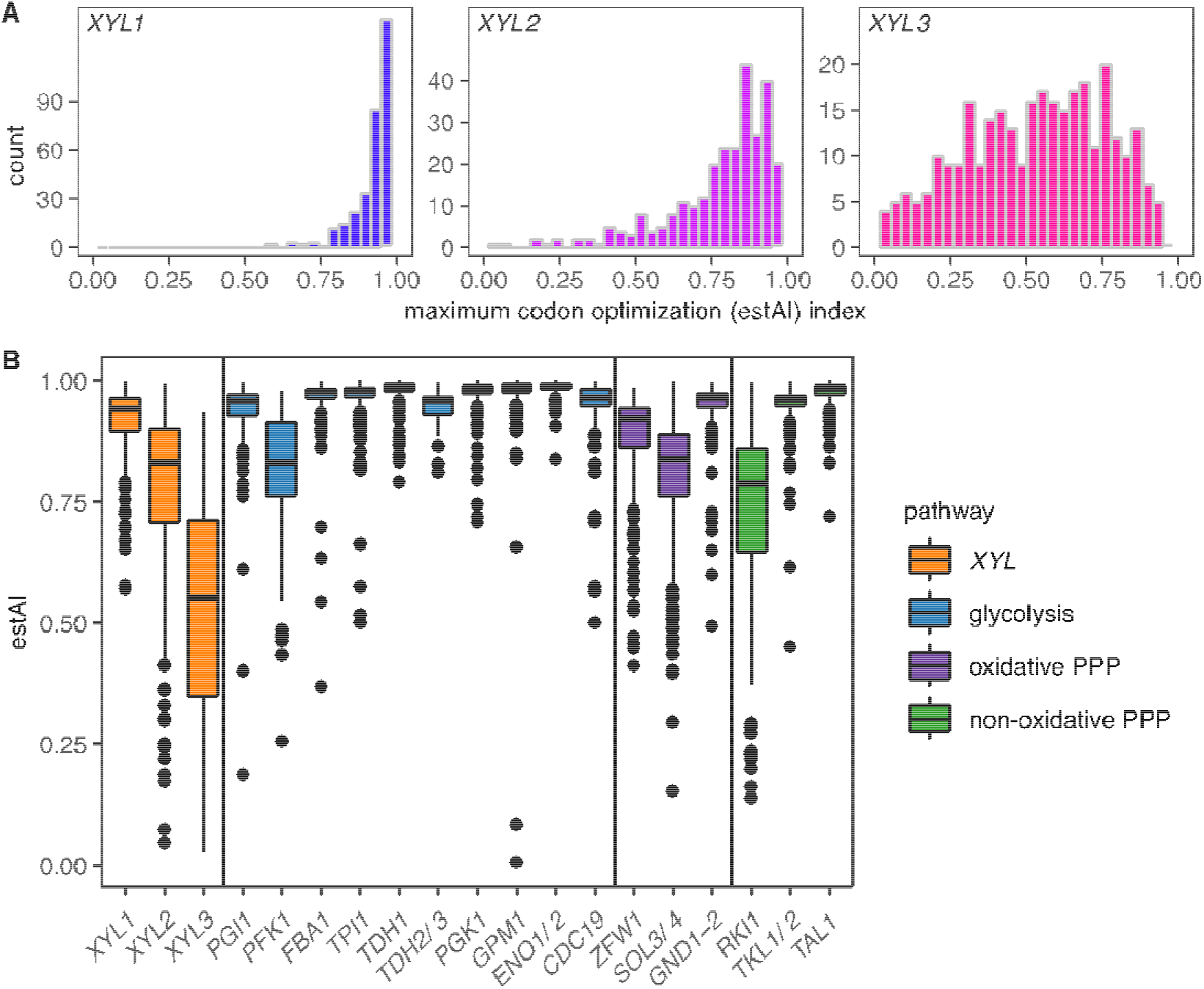
Distribution of codon optimization indices (estAI values). A) Histograms of the distribution of maximum estAI value among 320 of the 325 species for *XYL1, XYL2*, and *XYL3* are shown. *XYL1* genes are skewed towards highly optimized (blue), *XYL*2 estAI values are somewhat less skewed (violet), and *XYL3* estAI values are broadly distributed (magenta). Median estAI values of 0.94, 0.83, and 0.55 were calculated for *XYL1, XYL*2, and *XYL*3, respectively. B) *XYL* gene estAI distributions were compared to other carbon metabolism pathways related to xylose metabolism. The *XYL* pathway (orange), in general, is less optimized than glycolysis (blue) or either branch of the pentose phosphate pathway (purple/green). Specifically, the *XYL1* distribution is significantly lower than the estAI distributions of highly expressed glycolytic genes (*FBA1, TPI1, TDH1, PGK1, GPM1, ENO1/ENO2*), but is similar to *PGI1. XYL2* genes have estAI values similar to the rate-limiting steps in glycolysis (*PFK1*) and the oxidative pentose phosphate pathway (*ZWF1*). *XYL3* was less optimized on average than genes involved in glycolysis or the pentose phosphate pathway (PPP).

To provide context to codon optimization index distributions for *XYL* genes, we compared them to the optimization indices of genes that function in glycolysis and the pentose phosphate pathway (Fig. 3B). The *XYL1* distribution was lower than the estAI distributions of highly expressed glycolytic genes (*FBA1, TPI1, TDH1, PGK1, GPM1, ENO1/ENO2*), and was similar to *PGI1*, which encodes the glycolysis-initiating enzyme phosphoglucose isomerase. *XYL2* genes were less codon-optimized than most glycolytic genes, but interestingly, the *XYL2* estAI distribution was similar to the rate-limiting steps in glycolysis (*PFK1*) and the oxidative pentose phosphate pathway (*ZWF1*). *XYL3* was clearly less codon-optimized on average than genes involved in glycolysis or the pentose phosphate pathways.

### Codon optimization predicts xylose growth abilities

We next tested whether codon optimization of *XYL* genes was predictive of the ability to utilize xylose as a carbon source by comparing the estAI distributions of species that can grow on xylose to those that cannot for each gene. We limited our comparison to the 234 species with complete pathways and for which both estAI values and growth data were measured (Table S1). *XYL* gene codon optimization was significantly predictive of growth on xylose for all three genes examined (Wilcoxon rank sum test, *XYL1* p=8.9×10^−9^, *XYL2* p=3.7×10^−4^, *XYL3* p=9.5×10^−7^; Fig. 4A). This effect was not an artifact of phylogenetic correlation, as the trend was largely consistent across clades (Fig. S8).

**Figure 4.**
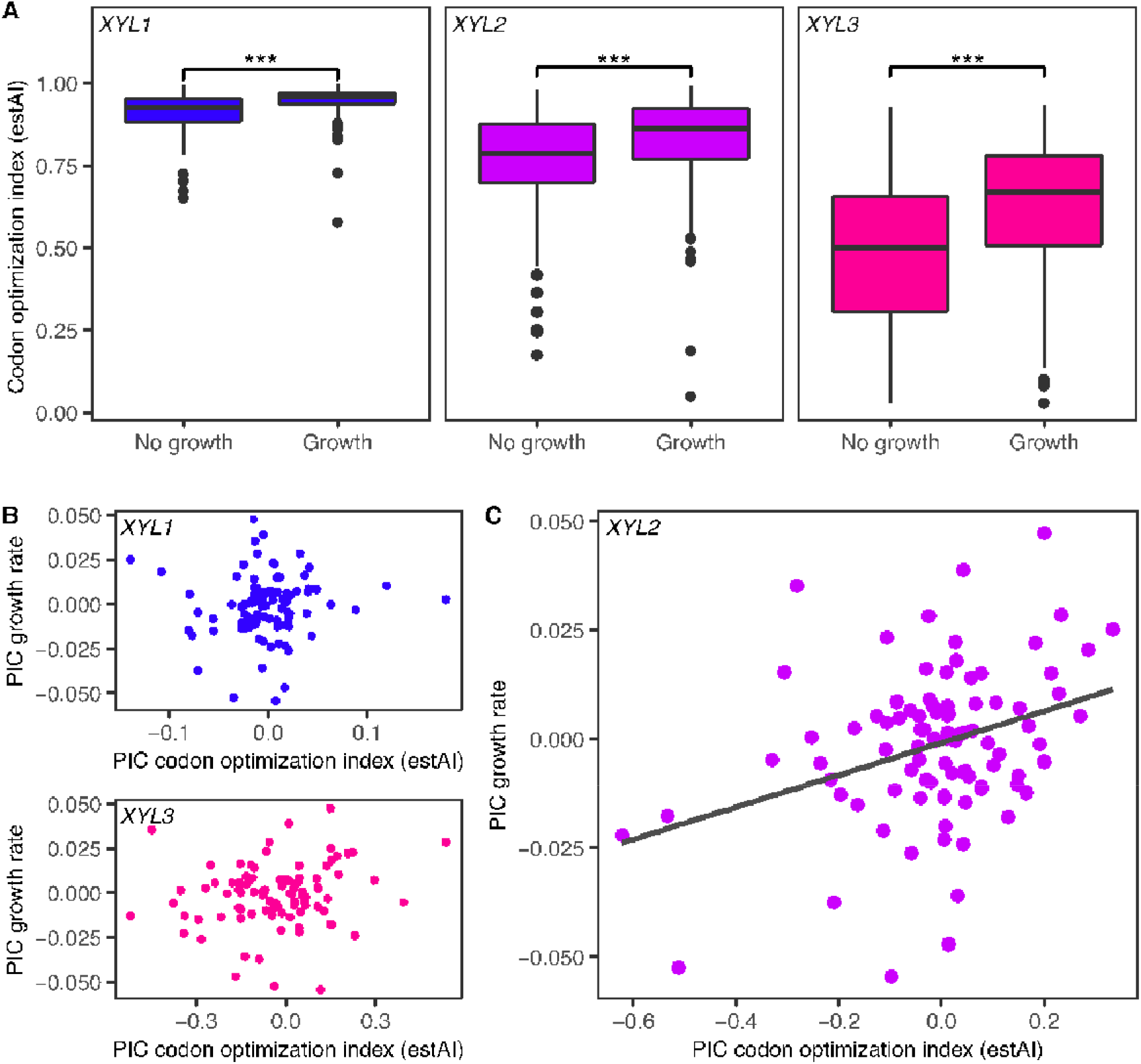
Species able to metabolize xylose have more highly codon-optimized *XYL* genes. A) Boxplots showing the distribution of estAI values for species unable to use xylose (left) compared to those that can (right) for *XYL1* (blue), *XYL2* (violet), and *XYL3* (magenta). (Wilcoxon rank sum test ***, *XYL1* p=8.9×10^−9^, *XYL2* p=3.7×10^−4^, *XYL3* p=9.5×10^−7^). This effect is not an artifact of phylogenetic correlation, as the trend is largely consistent across clades (Fig. S8). B-C) Phylogenetically independent contrast (PIC) analyses of *XYL1, XYL2*, and *XYL3* estAI in relation to xylose growth. *Kodamaea laetipori* and *Blastobotrys adeninivorans* were removed as outliers prior to analyses. Codon optimizations of *XYL1* and *XYL3* do not correlate with xylose growth rates. Codon optimization of *XYL2* is significantly correlated with growth rate in xylose medium (p=9×10^−4^, adj r^2^=0.11).

### Codon optimization of *XYL2* correlates with xylose growth rates

We have shown previously that codon optimization indices of specific genes involved in galactose metabolism not only predict whether a budding yeast species can utilize galactose, but can also be used to predict the rates of growth on galactose (33). We similarly compared *XYL* gene codon optimization to growth rates measured in medium containing xylose as the sole carbon source to determine whether this trait would be useful in predicting xylose metabolism as well. Phylogenetically independent contrasts (PICs) were used to compare estAI values and growth rates for the 93 species with complete pathways and for which there was previously published evidence of selection on codon usage (31). Of the three genes examined, only *XYL2* had a significant correlation between codon optimization and growth rate (p=9×10^−4^, adj r^2^=0.11; Fig. 4B-C).

## Discussion

Xylose fermentation is an ecologically important trait of immense biotechnological value for the conversion of sustainable plant feedstocks into biofuels. This study is the first to identify systematically *XYL* pathway homologs across a wide breadth of Saccharomycotina that includes all 12 major clades. While most genomes examined contain complete pathways, less than half of those species were able to assimilate xylose under laboratory conditions. In contrast, other metabolic traits that have been investigated in yeasts exhibit strong gene-trait associations (10,18). In particular, a survey of galactose metabolism across the same extensive collection of budding yeast species found that 89% of species with complete *GAL* pathways were able to use galactose as a carbon source (33). The unique inability of gene content to predict xylose-metabolism traits has been noted before (11,17,18), but not to this scale. Xylose metabolism patterns vary amongst yeast clades; most CUG-Ser1 species are able to utilize xylose, assimilation shows up sporadically in most other clades, and it is completely absent in the Saccharomycetaceae. These patterns are consistent with previous observations (reviewed in 39). To date, it has remained unclear why so many species with intact xylolytic genes appear nonetheless unable to use xylose. We find that, in part, this discrepancy can be explained by variation in translational selection on codon usage.

One limitation of this study and a possible explanation for the poor correlation between genotype and phenotype is that xylose catabolism only occurs in specific conditions. We analyzed only growth data generated in our assay under a single controlled condition. For some species, our data conflict with data aggregated from species descriptions (40). For other species, conflicting data also exist elsewhere in the literature. For example, *Kluyveromyces marxianus* did not grow in our 96-well plate assay but has been found to consume xylose in shake flasks (41). Oxygenation, base media, and temperature have all been documented as affecting xylose metabolism in different yeast species (3,42). Beyond condition dependence, intraspecific metabolic heterogeneity, such as known to occur in *Kluyveromyces lactis* and *Torulaspora delbrueckii*, could also produce inconsistencies (43,44). A final reason why our data may conflict with pre-existing descriptions is historical human errors in species typing and identification (45). Our choice to confine our analysis to the data we directly collected from taxonomic type strains may have obscured growth in a few species, but in general, it eliminated the effects of inconsistent conditions and taxonomical error.

We report here that metabolic gene codon optimization is useful for identifying species with xylolytic traits. Although the codon optimization index profiles differ markedly amongst the three *XYL* pathway genes, we find remarkable consistency in the difference in codon optimization between species that can and cannot metabolize xylose with a similar effect size for all three genes. Thus, codon optimization of individual *XYL* genes can be used to predict xylose metabolism from genome sequence data alone in budding yeasts. It is tempting to speculate that these data suggest that high *XYL* gene expression levels are necessary for xylose assimilation. Indeed, heterologous *XYL* gene expression levels have repeatedly been shown to be important determinants of xylose fermentation in engineered *S. cerevisiae* (46,47). However, higher codon optimization indices could also reflect directional selection on *XYL* gene codon usage in species for which xylose is an ecologically relevant sugar. Therefore, higher estAI values are likely observed because the species consumes xylose, rather than the inverse. This explanation does not preclude an effect of codon optimization on *XYL* expression, but it likely means that codon optimization is not sufficient for xylose utilization, even though it is highly predictive.

The *XYL1* and *XYL2* phylogenies we generated show evidence of widespread duplication and loss. Full elucidation of the gene history of these groups will require additional sampling both within and outside of the Saccharomycotina. However, differences between the species distributions of retained paralogs of *XYL1* and *XYL2* are curious. In the case of *XYL1*, most duplicated lineages are found in clades with high rates of xylose utilization. Detailed studies of *XYL1* paralog pairs within the CUG-Ser1 clade have found evidence of divergence in cofactor preferences (16,20), and *XYL1* duplications have generally been thought of as adaptations for xylose metabolism. However, our subphylum-wide analysis did not find a general relationship between *XYL1* copy number and xylose metabolism.

The consequences of *XYL2* duplication and the selective pressures driving *XYL2* duplicate retention have received far less attention. Given the seeming ecological irrelevance of xylose utilization in the Saccharomycetaceae, the diversification and retention of *XYL*2 genes in this group lacks a clear explanation unless the primary function of *XYL2* homologs in this family is not in xylose degradation. Several lines of evidence in the literature that support this notion: 1) there is ample evidence that budding yeast XDH enzymes are promiscuous across polyols (48– 51), 2) the *XYL2* reverse reaction (reduction of xylulose to xylitol) is more energetically favorable by an order of magnitude (52), and 3) the strongest phylogenetic signal of *XYL* gene loss we observed was in the W/S clade of yeasts, which is a group of fructose-specializing yeasts that have evolved a novel means of reducing fructose to maintain redox balance (53). Taken together, these data are suggestive of an alternative role of the *XYL* pathway, and *XYL2* in particular. Instead of supporting xylose utilization, XDH activity in these yeasts may be important for regenerating oxidized NAD^+^ in certain growth conditions through the reduction of sugars, including xylulose, fructose, and mannose, to the polyols xylitol, sorbitol, and mannitol, respectively. Additional experimental work in the family Saccharomycetaceae is needed to determine if XDH activity plays a role in redox balance as suggested above, or perhaps functions in a yet-to-be-discovered process.

We also find *XYL2* is the only *XYL* gene for which codon optimization has a linear relationship with growth rate on xylose. *XYL2* has repeatedly been implicated as the rate-limiting step in xylose metabolism (54–56). The correlation between codon optimization and growth that we report supports the hypothesis that endogenous *XYL2* expression levels affect rates of xylose consumption in natively xylose-consuming yeasts. This optimization could be, in part, to overcome the unfavorable reaction kinetics and subpar substrate specificity mentioned above. Interestingly, the *XYL2* estAI distribution we observed was highly similar to that of rate-limiting steps of glycolysis (*PFK1*) and the oxidative pentose phosphate pathway (*ZWF1*), perhaps pointing to a general trend in genes encoding enzymes with rate-limiting or regulatory roles.

Xylose metabolism cannot be predicted by gene content in budding yeasts. Here, we show that there is a significant predictive value of codon optimization in the detection of native xylose-metabolizing yeasts across all three genes required for xylose degradation. Xylose fermentation is a trait of great ecological and biotechnological interest, while being exceedingly rare. Instead of expending resources testing large sets of yeasts or their synthesized genes, codon optimization could be used to filter for candidate yeasts with a higher probability of containing highly xylolytic pathways. We also show that *XYL2* optimization has a linear relationship with growth rates on xylose. In the absence of growth or metabolic data, *XYL2* sequences can be used to predict which species are likely to catabolize xylose especially well. This work presents a novel framework of leveraging signatures of selection, specifically codon optimization, for understanding weak and variable gene-trait associations and could be a valuable tool for understanding trait variation in other systems.

## Materials and Methods

### Identification of *XYL1, XYL2*, and *XYL3* homologs

We identified homologs of *XYL1, XYL2*, and *XYL3* across 332 published budding yeast genome assemblies (10) using Hidden Markov Model (HMMER) sequence similarity searches (v3.3 http://hmmer.org). HMM profiles were built using sequences retrieved from a BLASTp search using *Spathaspora passalidarum XYL1*.*1, XYL2*.*1*, and *XYL3*. Hits were manually curated to retain an alignment of fourteen sequences representing a phylogenetically diverse taxon set. HMMER searches were performed on protein annotations generated with ORFfinder (NCBI RRID:SCR_016643) using default settings, which include nonconventional start codons. Sequences were later manually curated to confirm true start sites (see below). We did not account for modified translation tables found in some yeast clades (CUG-Ser1, CUG-Ser2, and CUG-Ala clades (10)) because this codon is known to be rare (31).

HMMER searches for *XYL1* and *XYL2* both identified large gene families of aldose reductases and medium-chain dehydrogenases, respectively. To identify the *XYL* orthologous sequences, HMMER hits were assigned KEGG orthology with BLASTKoala (57), and approximate maximum likelihood trees of KEGG-annotated hits were built with FastTree v2.1.10 (58) (Fig. S1-S2). Subclades containing *XYL* gene homologs based on KEGG orthology (*XYL1* - K17743, *XYL2* - K05351) were identified for *XYL1* and *XYL2*.

Coding sequences of homologs for all three genes were then manually curated. True start sites were identified using TranslatorX (59), and sequences were trimmed or expanded accordingly. A combination of alignment visualization and collapsed tree inspection was used to identify highly divergent sequences that were then examined via BLAST; likely bacterial contaminants were removed. Maximum likelihood phylogenies of protein sequences for each of the three genes were built with IQTree (60) using ModelFinder automated model selection (61) (*XYL1*-LG+F+I+G4, *XYL2-*LG+I+G4, *XYL3*-LG+F+I+G4) based on 1,000 bootstrap replications. An independent maximum likelihood tree of *XYL2* protein sequences in the family Saccharomycetaceae with the addition of *S. cerevisiae XDH1* (37) was generated using IQ tree with an LG+I+G4 substitution model and node support based on 1,000 bootstrap replications. Trees were visualized and annotated in iTOL (62).

### Growth assays

All yeast strains used in growth experiments were first plated on Yeast Extract Peptone Dextrose (YPD) agar plates and grown until single colonies were visible. The plates were then stored at 4°C for up to a month. Single colonies were then cultured in liquid YPD for a week at room temperature on a culture wheel. After a week of growth, yeast strains were subcultured in 96-well plates containing Minimal Medium with 1% glucose or 1% xylose and allowed to grow for a week at room temperature. The 96-well plates contained a 4 quadrant moat around the edge of the plate where 2mL of water was added to each quadrant. The addition of water to the plate prevents evaporation in the edge and corner wells, allowing for the whole plate to be utilized. After the initial week of growth on the treatments, all yeasts were transferred into fresh 1% glucose or 1% xylose minimal medium and placed on a plate reader and stacker (BMG FLUOstar Omega). Plates were read every two hours for a week at OD600. All growth experiments were replicated three times. In each replicate, both the order of yeasts on the plate and order of sugars on the plate were randomized to alleviate plate effects. Growth rates were quantified in R using the package *grofit* (63). Average growth rates were calculated across replicates for each species.

### Codon Optimization

Codon optimization indices of *XYL1, XYL2*, and *XYL3* homologs were determined as in LaBella et al. (33). Briefly, species-specific codon optimization values (wi values) retrieved from (31) were used to calculate the codon optimization index (stAI) for each ortholog identified by calculating the geometric mean of species-specific wi values for each gene. Five species in our dataset do not have corresponding wi values and were dropped from codon optimization analyses (*Middelhovenomyces tepae, Nadsonia fulvescens* var. *fulvescens, Spencermartinsiella europaea, Botryozyma nematodophila*, and *Martiniozyma abiesophila*). To compare codon optimization values between species, we normalized gene-specific stAI values to the genome-wide distribution of stAI values for each species using the empirical cumulative distribution function. The resulting normalized codon optimization index (estAI value) is an estimate of the genome-wide percentile of codon optimization for each gene (e.g. an estAI value of 0.95 indicates a gene that is more optimized than 95% of genes in the genome). For species with multiple paralogs, including those derived from the whole genome duplication, only the gene with the highest estAI value was considered in further analysis.

Orthologs of glycolysis pathway genes (*CDC19, ENO1/ENO2, FBA1, GPM1, PFK1, PGI1, PGK1, TDH1, TDH2/TDH3, TPI1*) and pentose phosphate pathway genes (*GND1/GND2, RKI1, SOL3/SOL4, TAL1, TKL1/TKL2, ZWF1*) were identified using HMMER searches as described above with the exception of manual curation. Codon optimization for each gene was measured as described above. For species with multiple paralogs, only the maximum estAI value per gene per species was retained for analysis.

### Statistical Analyses of Growth Data and Codon Optimization

We compared codon optimization (estAI) values of species with non-zero growth rates in xylose medium to species that did not grow using a two-sample Wilcoxon test for all three genes. To compare xylose growth rates to estAI values, we first retained data for only those species previously found to have evidence of genome-wide selection on codon usage (31). Two species had extremely high growth rates that did not appear to be artifactual (Fig. S3). Since phylogenetic independent contrasts are highly sensitive to outlier data, we removed these two species. For the remaining 93 species, growth rate was compared to codon optimization by fitting a linear model to phylogenetically independent contrast (PIC) values to account for phylogenetic relatedness. PIC values were generated using the ape package in R (64). All other statistical analyses were performed using R stats v3.6.2.

## Acknowledgments

We thank members of the Hittinger and Rokas groups for helpful discussions. This material is based upon work supported by the National Science Foundation under Grant Nos. DEB-1442148, DEB-2110403, DEB-1442113, and DEB-2110404; in part by the DOE Great Lakes Bioenergy Research Center (DOE BER Office of Science DE-SC0018409); and the USDA National Institute of Food and Agriculture (Hatch Project 1020204). CTH is an H. I. Romnes Faculty Fellow, supported by the Office of the Vice Chancellor for Research and Graduate Education with funding from the Wisconsin Alumni Research Foundation. Research in AR’s lab is also supported by the National Institutes of Health/National Institute of Allergy and Infectious Diseases (R56 AI146096 and R01 AI153356), and the Burroughs Wellcome Fund. KJF is a Morgridge Metabolism Interdisciplinary Fellow supported by the Morgridge Institute for Research -Metabolism Theme.

## Data availability

Analyses were performed on the 332 published and publicly available assemblies analyzed in Shen et al. 2018. Codon optimization values were obtained from the figshare repository from LaBella et al. 2019 (https://doi.org/10.6084/m9.figshare.c.4498292). All data generated in this project, including *XYL* gene sequences, may be requested from the corresponding author and will be publicly archived upon publication. The raw data used to generate each figure is reported in data file S1.

## Supplemental data files

S1. Excel file containing all raw data associated with figures.

## Notes

### Competing Interest Statement

A.R. is a scientific consultant for LifeMine Therapeutics, Inc.

