## Supplementary File for "Codon optimization, not gene content, predicts *XYL*ose metabolism in budding yeasts"

Supplemental Material

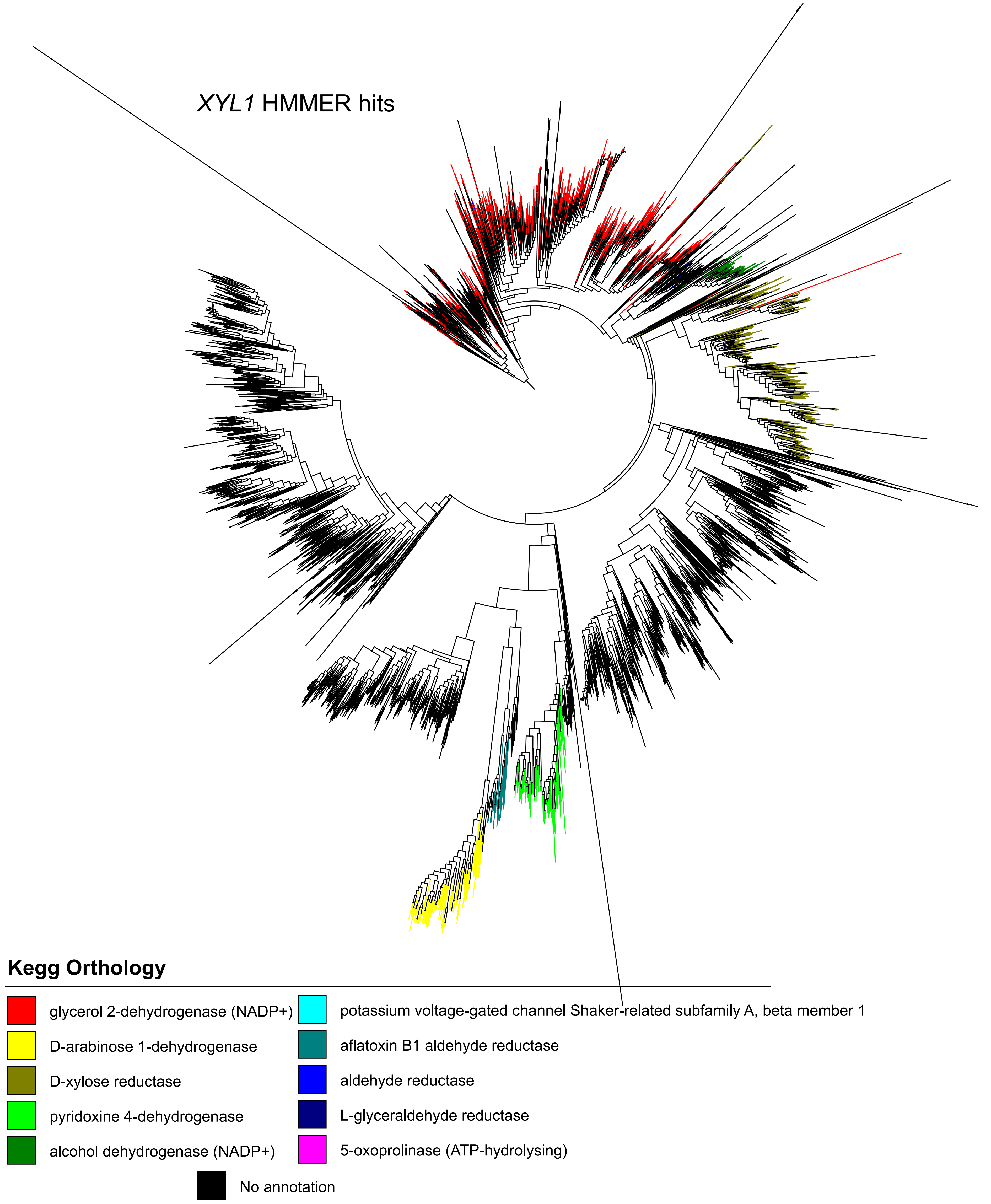


**S1. *XYL1* homologs identified in sequence similarity search.** HMMER search for *XYL1* homologs identified a large gene family of aldo-keto reductases. Genes were annotated by KEGG orthology via BLASTKoala. Maximum likelihood tree of *XYL1* KEGG-annotated hits were built with FastTree. The subclade containing xylose reductase homologs was identified for manual sequence curation (dark tan, *XYL1* - K17743).

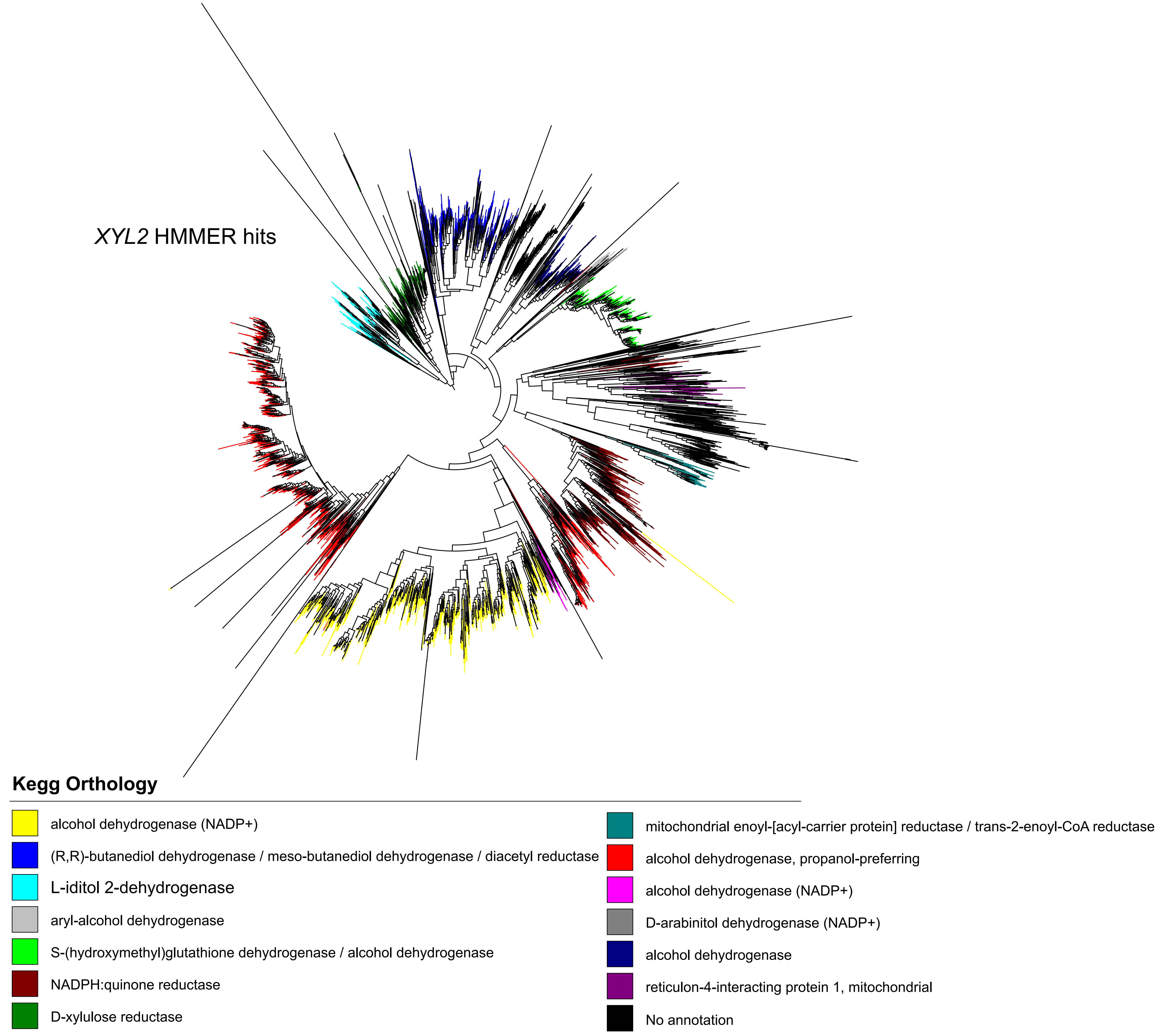


**S2. *XYL2* homologs identified in sequence similarity search.** HMMER search for *XYL2* homologs identified a large gene family of medium-chain dehydrogenases. Genes were annotated by KEGG orthology via BLASTKoala. Maximum likelihood tree of *XYL2* KEGG-annotated hits were built with FastTree. The subclade containing xylitol dehydrogenase (xylulose reductase) homologs was identified for manual sequence curation (dark green, *XYL2* - K05351).

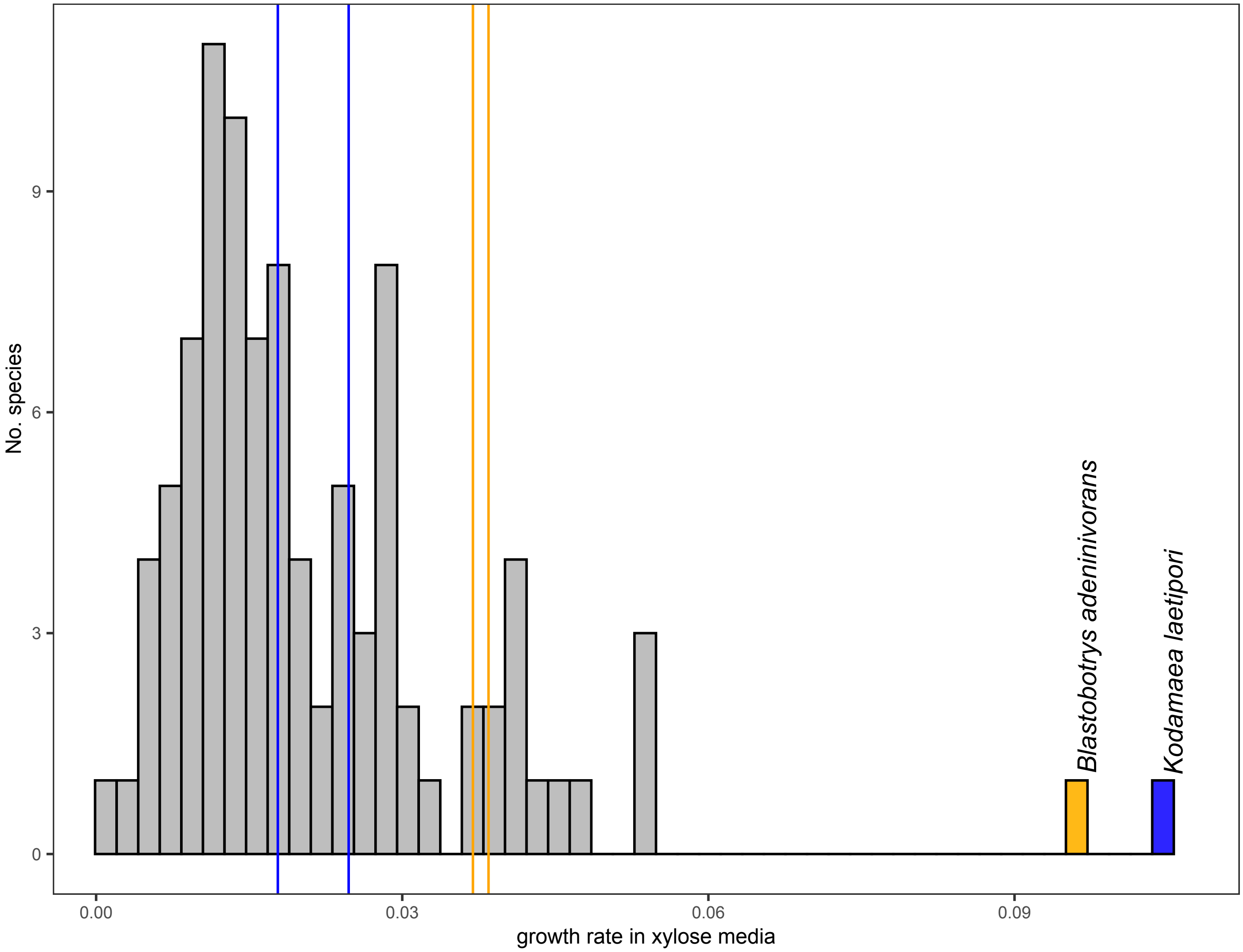


**S3. Distribution of xylose growth rates.** Growth rates in medium containing xylose as a sole carbon source were measured for 281 species. *Kodamaea laetipori* (blue) and *Blastobotrys adeninivorans* (orange) both have much higher growth rates than their closest relatives (blue lines - *Candida parapsilosis* and *Candida restingae*, orange lines - *Blastobotrys raffinosifermentans* and *Blastobotrys peoriensis*). *Kodamaea laetipori* and *Blastobotrys adeninivorans* were removed as outliers prior to generating phylogenetically independent contrasts for growth rate data.

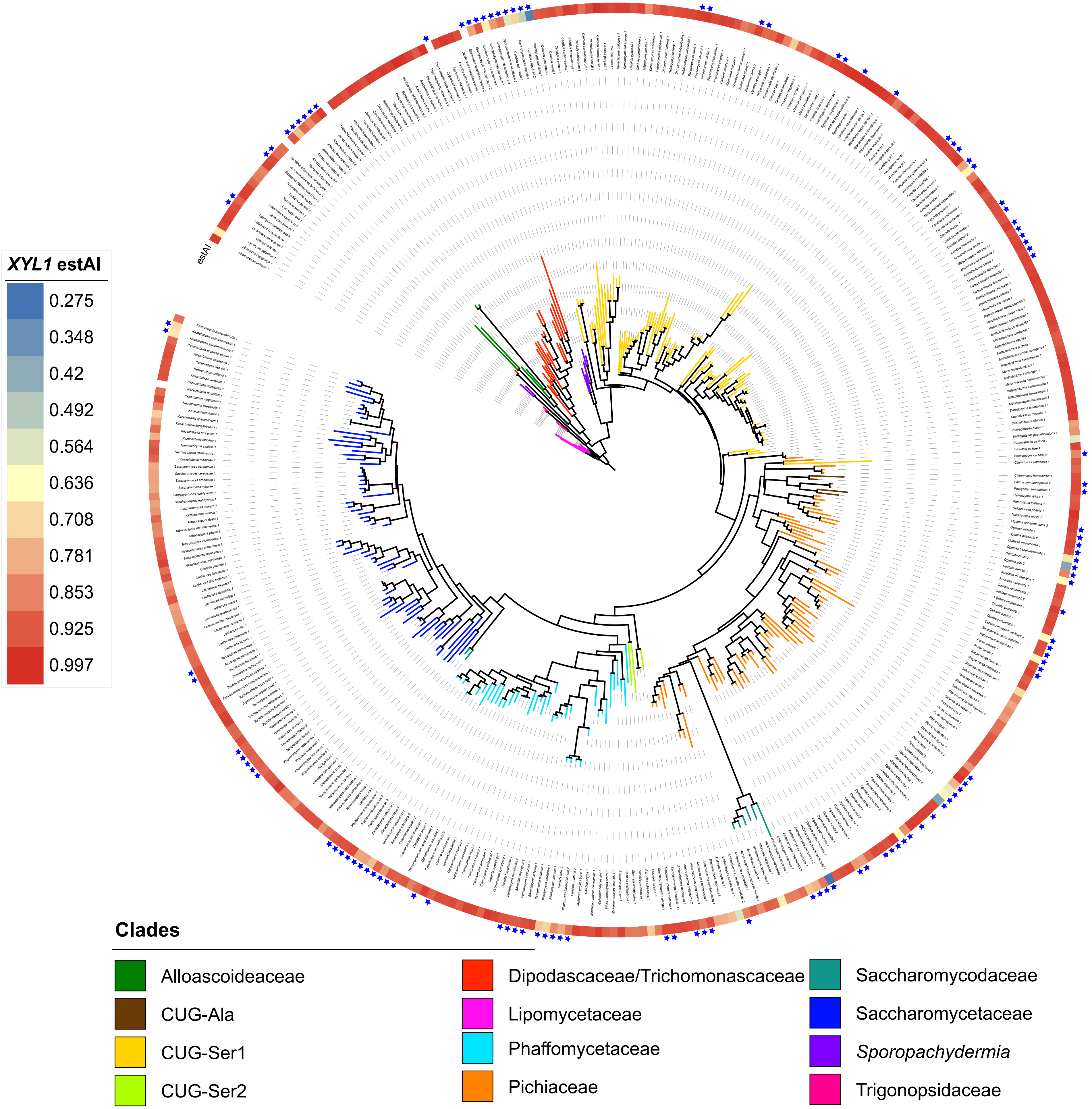


**S4. Maximum likelihood tree of Xyl1 protein sequences.** Maximum likelihood phylogeny of Xyl1 protein sequences was built using IQTree with an Auto substitution model based on 1,000 bootstrap replications. Major yeast clades are depicted by branch color using tree topology from Shen et al. (1). Color gradient indicates codon optimization indices (estAI values) with color ranging from blue (lowest estAI value) to red (highest estAI value). Stars indicate paralogs belonging to multi-copy lineages. Note that these analyses show that *S. cerevisiae GRE3* is a *XYL1* homolog.

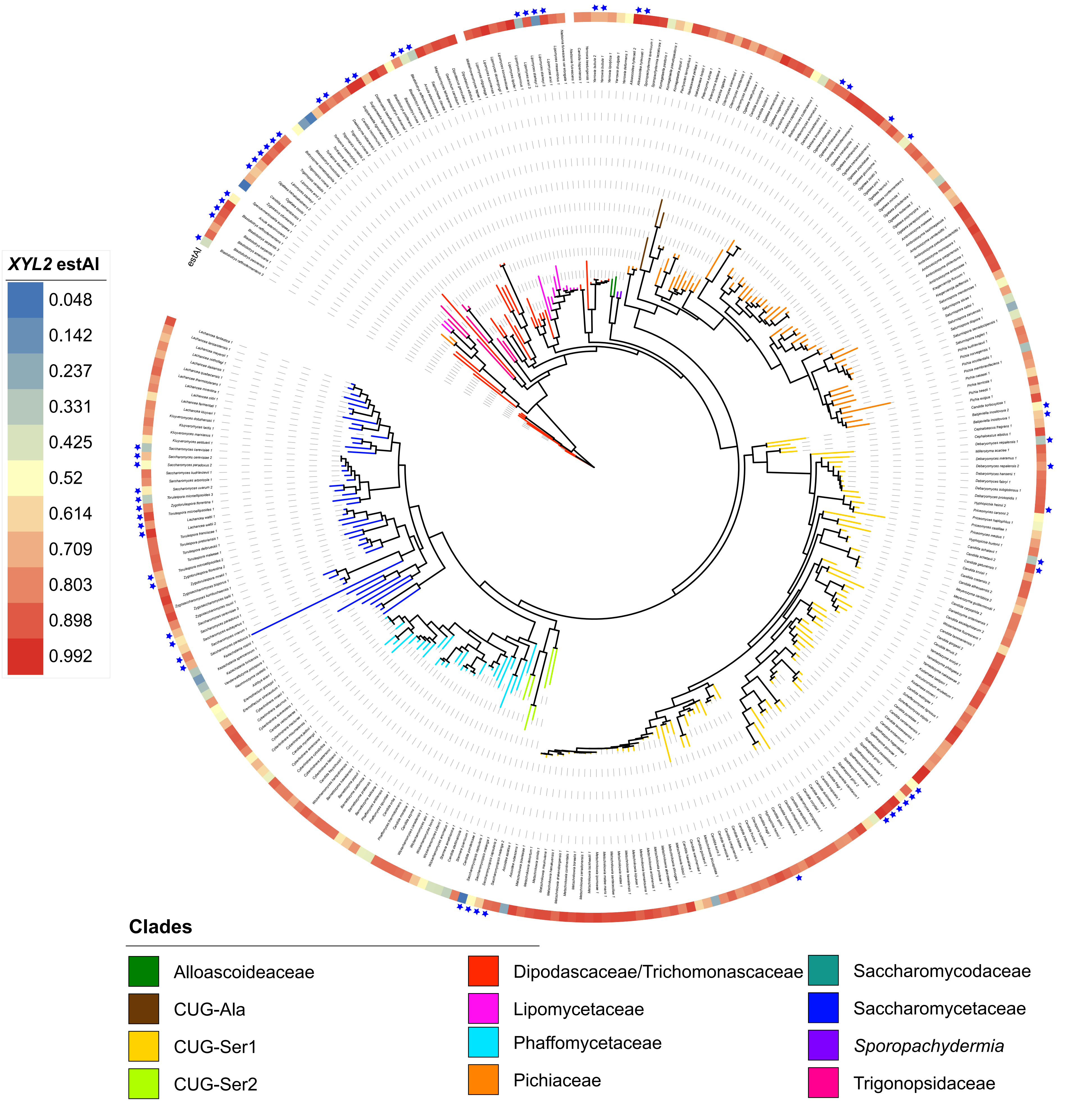


**S5. Maximum likelihood tree of Xyl2 protein sequences.** Maximum likelihood phylogeny of Xyl2 protein sequences was built using IQTree with an Auto substitution model based on 1,000 bootstrap replications. Major yeast clades are depicted by branch color using tree topology from Shen et al. (1). Color gradient indicates codon optimization indices (estAI values) with color ranging from blue (lowest estAI value) to red (highest estAI value). Stars indicate paralogs belonging to multi-copy lineages. Notably, the stars depict the lineage-specific duplication of *SOR1*/*SOR2* in *S. cerevisiae*. The nearly identical *S. cerevisiae* paralogs *SOR1* and *SOR2* fall within the Saccharomycetaceae *XYL2* clade, suggesting that *SOR1*/*SOR2* are both paralogs of S. cerevisiae *XYL2*. Associated *S. cerevisiae* gene nomenclature: *Saccharomyces cerevisiae 1 - SOR2, Saccharomyces cerevisiae 2 - SOR1, Saccharomyces cerevisiae 3 - XYL2*.

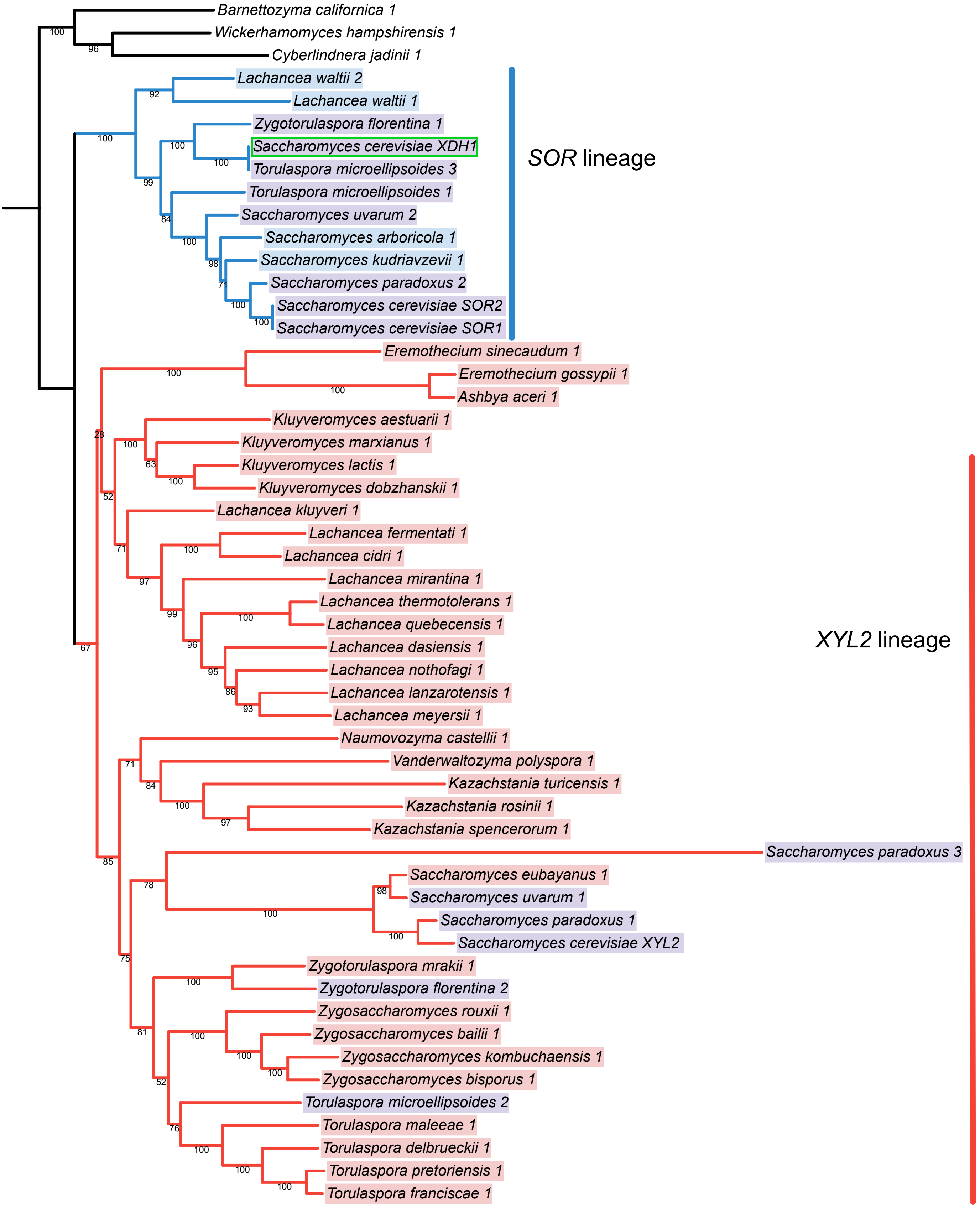


**S6. Maximum likelihood tree of Saccharomycetaceae Xyl2 protein sequences.** Independent maximum likelihood tree of Xyl2 protein sequences in the family Saccharomycetaceae with the addition of *S. cerevisiae XDH1* (2) (green box) generated using IQ tree with an LG+I+G4 substitution model and node support based on 1,000 bootstrap replications. The duplication of *XYL2* prior to the yeast whole genome duplication resulted in two major lineages sorting within the Saccharomycetaceae, the *SOR* lineage (blue) and the *XYL2* lineage (red). Species that retained sequences from both lineages are indicated in purple. *XDH1* (green box), previously identified as responsible for weak xylose utilization in some wine *S. cerevisiae* strains, was included and found to be horizontally transferred from *Torulaspora microellipsoides* (3).

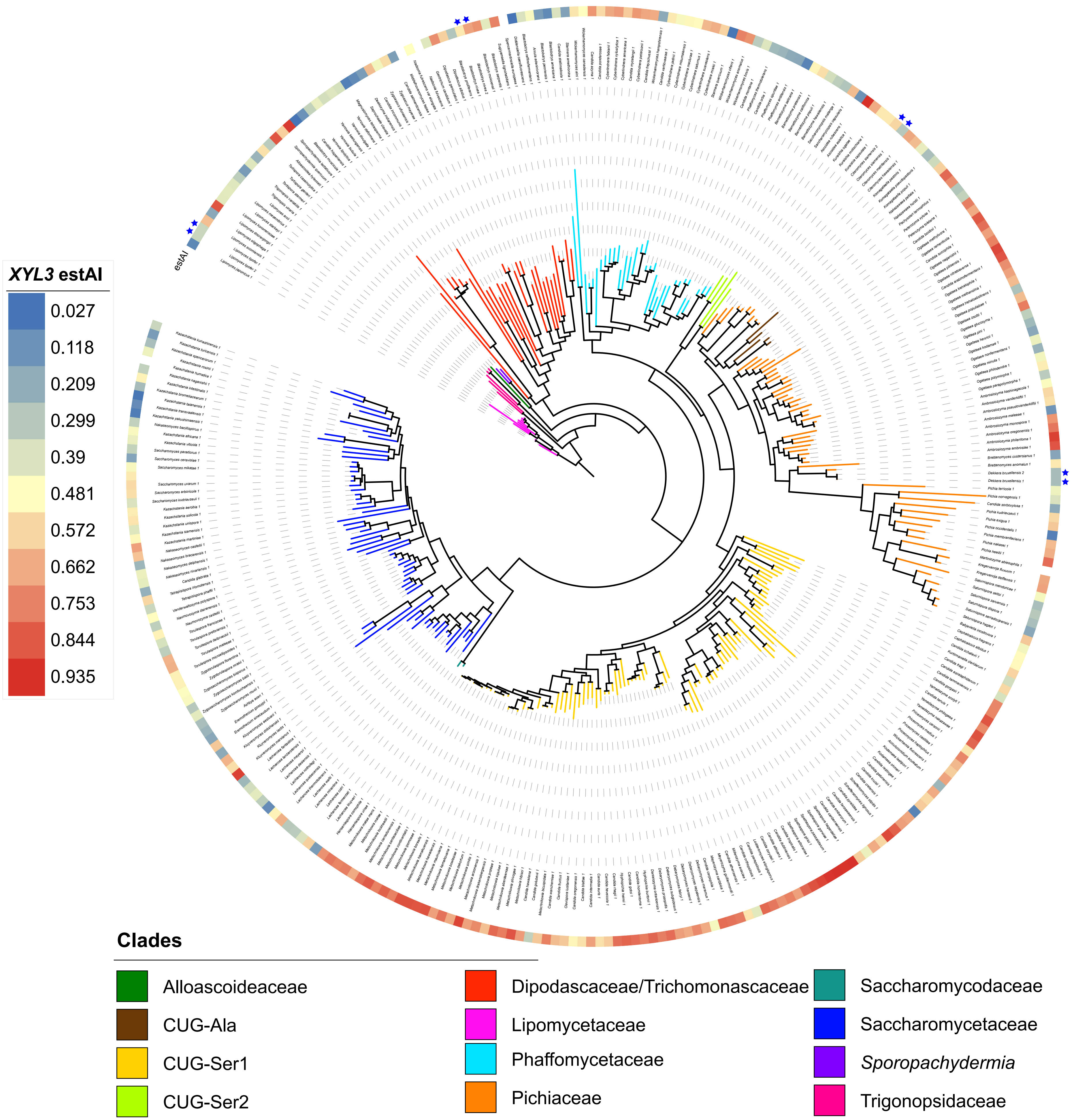


**S7. Maximum likelihood tree of Xyl3 protein sequences.** Maximum likelihood phylogeny of Xyl3 protein sequences was built using IQTree with an Auto substitution model based on 1,000 bootstrap replications. Major yeast clades are depicted by branch color using tree topology from Shen et al. (1). Color gradient indicates codon optimization indices (estAI values) with color ranging from blue (lowest estAI value) to red (highest estAI value). Stars indicate paralogs belonging to multi-copy lineages.

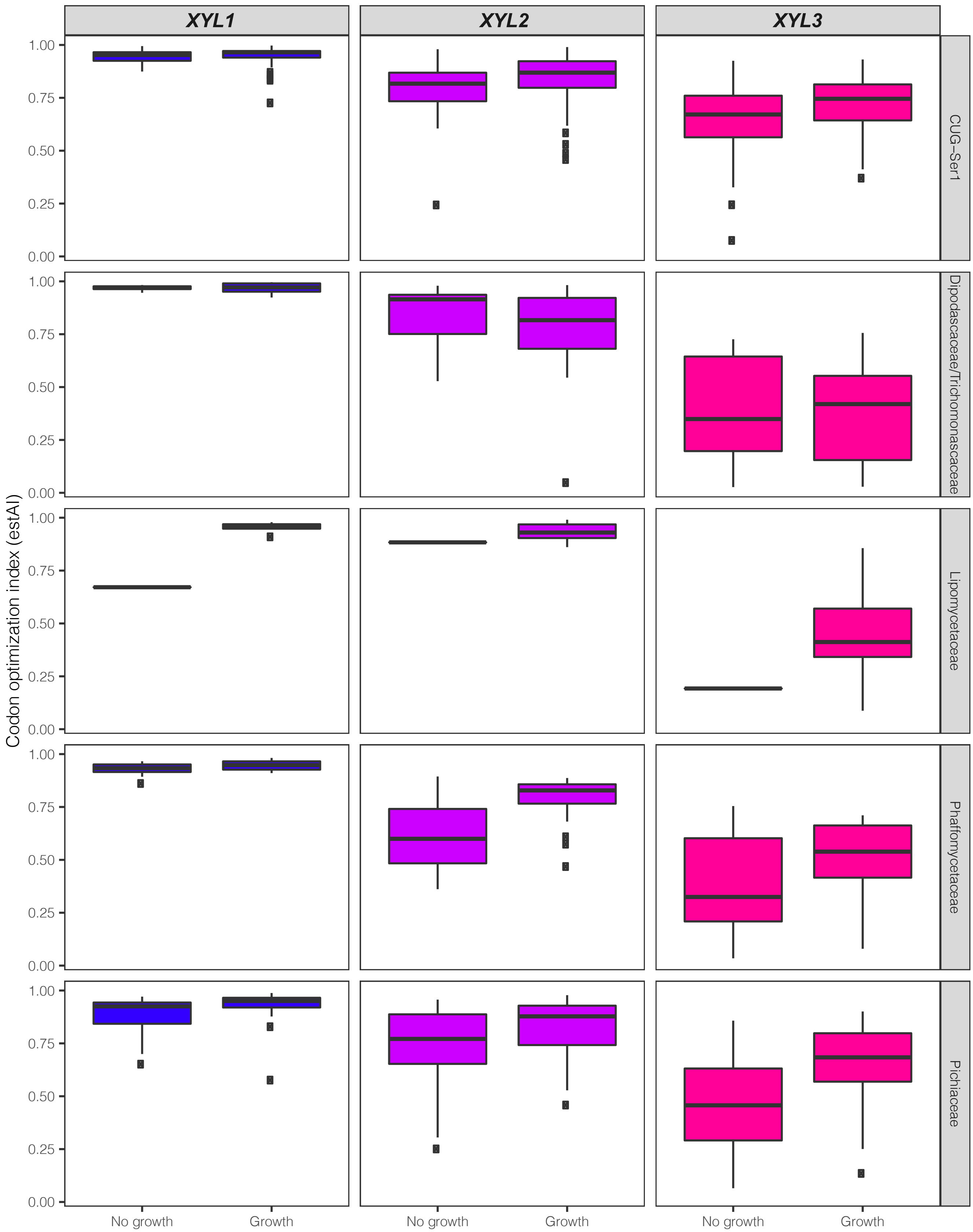


**S8. Comparison of codon optimization indices with xylose growth across clades.** Codon optimization was plotted for each of the 5 clades that contain growers, non-growers, and more than 5 species. The trend that *XYL* gene codon optimization is significantly predictive of growth is largely consistent across clades. The consistency across clades supports the conclusion that this trend is not an artifact of phylogenetic correlation.
